# Serine 165 Phosphorylation of SHARPIN regulates the Activation of NF-κB

**DOI:** 10.1101/2020.07.02.184085

**Authors:** An Thys, Kilian Trillet, Sara Rosinska, Audrey Gayraud, Tiphaine Douanne, Yannic Danger, Clotilde Renaud, Luc Antigny, Régis Lavigne, Charles Pineau, Emmanuelle Com, Franck Vérité, Julie Gavard, Nicolas Bidère

## Abstract

The adaptor SHARPIN composes, together with the E3 ligases HOIP and HOIL1, the linear ubiquitin chain assembly complex. This enzymatic complex catalyzes and stamps atypical linear ubiquitin chains onto substrates to modify their fate, and has been linked to the regulation of the NF-κB pathway downstream most immunoreceptors, inflammation and cell death. However, how this signaling complex is regulated is not fully understood. Here, we report that a portion of SHARPIN is constitutively phosphorylated on the serine in position 165 in lymphoblastoid cells, and can be further induced following antigen receptor stimulation. Analysis of a phosphorylation-resistant mutant of SHARPIN revealed that this mark is dispensable for the generation of linear ubiquitin chains. However, phosphorylation allows the optimal activation of NF-κB in response to TNFα and T-cell receptor engagement. These results identify a new layer of regulation of the LUBAC, and unveil new strategies to modulate its action.

## Introduction

The linear ubiquitin chain assembly complex (LUBAC) is an enzymatic triad of the adaptor SHARPIN (SHANK-associated RH domain-interacting protein) and the E3 ligases HOIP (HOIL-1 interacting protein, also known as RNF31) and HOIL-1 (RanBP-type and C3HC4-type zinc finger-containing protein 1, also called RBCK1, HOIL-1L) [1–3]. This unique complex catalyzes the formation and attachments of atypical linear ubiquitin chains on substrates, thereby modifying their fate. The LUBAC acts as a linchpin by transducing signals from most immunoreceptors to NF-κB, and therefore emerges as a key regulator of innate and adaptive immunity [1–13]. For instance, the binding of tumor necrosis factor α (TNFα) to its cognate receptor TNFR1 (TNF receptor 1) drives the recruitment of the LUBAC into the so-called complex I. By decorating the key protein kinase RIPK1 (receptor-interacting protein 1 kinase) with linear ubiquitin chains, the LUBAC stabilizes and favors downstream NF-κB signaling [1–3,7]. Mice deficient in SHARPIN, HOIP or HOIL-1 are hallmarked by an exacerbated TNFα induced cell death [1–3,14–16]. The loss of LUBAC components destabilizes this TNFR signaling complex I and induces the assembly of cytosolic complex II, causing cell death by apoptosis or necroptosis [1,14,15]. While, mice carrying a spontaneous SHARPIN-null mutation (*cpdm*) develop multi-organ inflammation and chronic proliferative dermatitis HOIP and HOIL-1 deficiency is embryonically lethal [1–3,14–16]. The LUBAC also functions downstream of antigen receptors to ensure the optimal activation of NF-κB, and this signaling pathway is pirated in the activated B cell like subtype of diffuse large B-cell lymphomas (ABC DLBCL) [17–20]. Accordingly, the targeting of the LUBAC was shown to be toxic in ABC DLBCL, unveiling a contribution of this complex to lymphomagenesis [19–22].

How the LUBAC is regulated continues to be elucidated. All members of the LUBAC have been reported to carry linear ubiquitin chains through auto-ubiquitination [23,24]. Recently, Iwai and colleagues demonstrated that HOIL-1 E3 ligase mono-ubiquitinates the LUBAC, which causes HOIP to preferentially decorate the LUBAC with linear ubiquitin chains rather than other substrates [25]. Linear auto-ubiquitination of the LUBAC can be removed by the OTU deubiquitinase with linear linkage specificity (OTULIN) [23,24]. In addition, HOIP is processed upon TNFα- and TRAIL-induced apoptosis by caspases, with cleaved fragments displaying reduced NF-κB activation capabilities [26,27]. HOIP is also phosphorylated by mammalian ste20-like kinase 1 (MST1) in response to TNFα, and this modulates its E3 ligase activity, thereby attenuating NF-κB signaling [28]. Three independent groups, including ours, showed that HOIL-1 is cleaved by the paracaspase MALT1 upon antigen receptor engagement and constitutively in ABC DLBCL to allow optimal activation of NF-κB [29–31]. Lastly, SHARPIN is decorated with lysine (K) 63 ubiquitin chains in mice on K312. This ubiquitination was shown to be important for development of regulatory T cells [32]. However, what effect K63 ubiquitination of SHARPIN has on NF-κB signaling still remains an open question. Here, we demonstrate that a fraction of the LUBAC subunit SHARPIN is constitutively phosphorylated in lymphoblastoid cells, and this post-translational modification can be further induced upon antigen receptor engagement. We identify serine (S) 165 to be the primary phosphorylation site of SHARPIN, and provide evidence of its crucial role for the optimal activation of NF-κB response to both T-cell receptor engagement and TNFα stimulation.

## Results and discussion

### SHARPIN is a phosphoprotein

Western blotting analysis of SHARPIN, in human primary CD4^+^ and CD8^+^ cells, Jurkat cells and DLBCL cell lines, revealed that SHARPIN resolves as a doublet (Fig. 1A, EV1A). The treatment of cell lysates with lambda phosphatase, which removes phosphate groups from serine, threonine and tyrosine, effectively chased away the slow migration specie of SHARPIN, suggesting phosphorylation (Fig.1A, EV1A). Conversely, incubating cells with the phosphatase inhibitor Calyculin A resulted in an increase intensity of the slow migration specie of SHARPIN, reinforcing the idea that SHARPIN is indeed phosphorylated (Fig. 1A). Quantification of constitutive SHARPIN phosphorylation in numerous cell lines, revealed that phospho-SHARPIN exists at various levels within different cell types, with lymphoid cells having a relatively high SHARPIN phosphorylation (Fig. 1B). Hence, a fraction of SHARPIN is constitutively phosphorylated.

**Figure 1.**
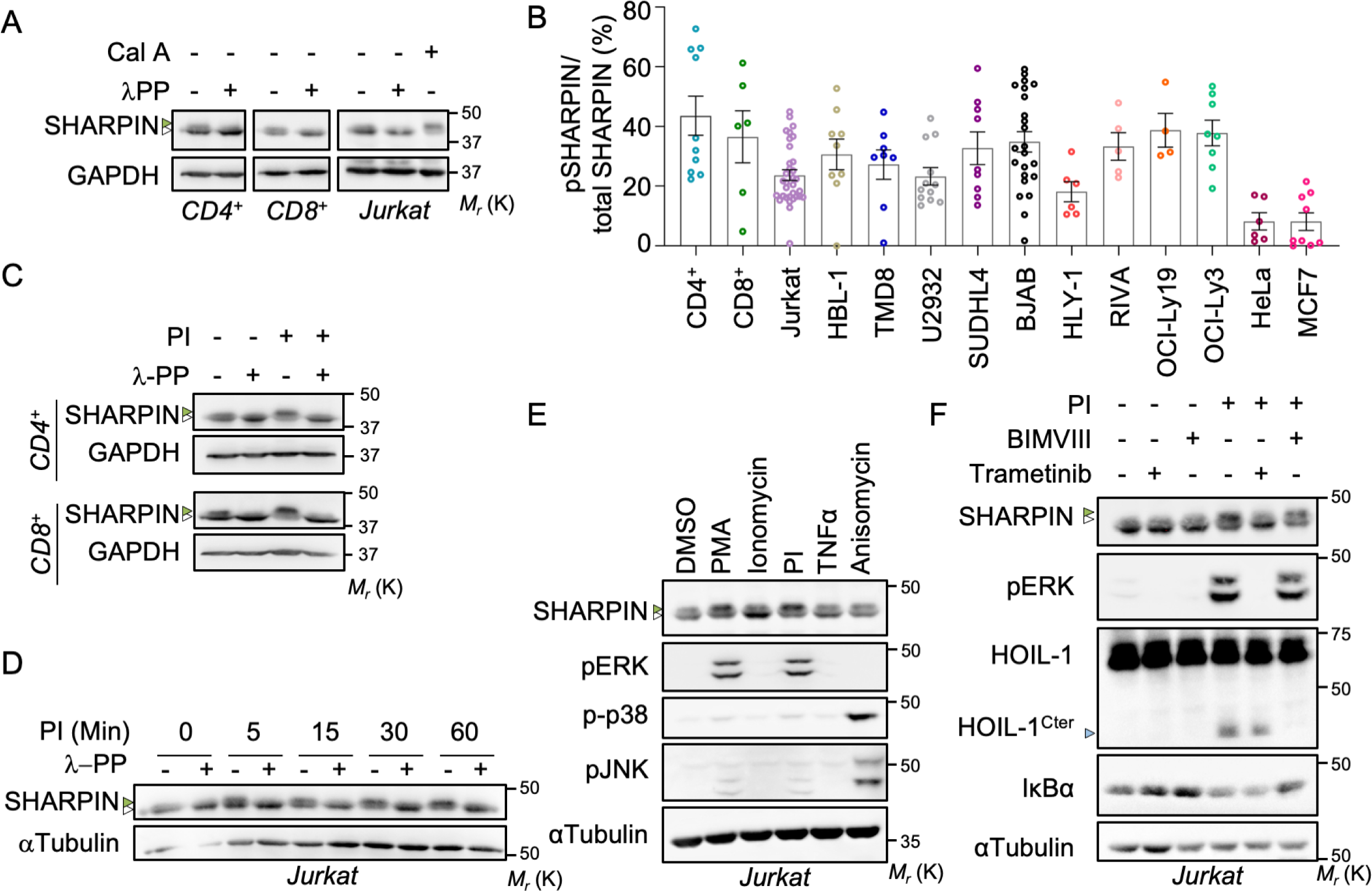
SHARPIN is a Phosphoprotein. A Jurkat cells were treated with or without 50 nM Calyculin A (Cal A) for 30 min prior lysis. Cell lysates from primary human CD4^+^ or CD8^+^ T lymphocytes and Jurkat cells were treated with lambda phosphatase (λPP) as indicated, and subjected to Western blotting analysis. Green and white arrowheads indicate phosphorylated and unphosphorylated SHARPIN, respectively. Molecular weight markers (*M*_*r*_) are indicated. B Densitometric analysis of the ratio between phosphorylated species of SHARPIN and SHARPIN in primary human T CD4^+^ cells, primary human T CD8^+^ T cells, Jurkat T cells, a panel of patient-derived diffuse large B-cell lymphoma cell lines, HeLa and MCF-7 cells. C-D Primary human CD4^+^ T (E) and CD8^+^ T lymphocytes (F), and Jurkat cells (G) were stimulated for 30 min or as indicated with 20 ng/mL PMA plus 300 ng/mL ionomycin (PI). Lysates were prepared and treated or not with λPP and Western blotting analysis was performed as indicated. E Jurkat cells were stimulated with 20 ng/mL PMA, 300 ng/mL ionomycin, PMA plus ionomycin (PI), 10 ng/mL TNFα or 12.5 µg/mL Anisomycin, for 30 min. Cell lysates were subjected to Western blotting analysis. F Jurkat cells we pretreated for 1h with 1 µM Trametinib or 500 nM BIMVIII and subsequently stimulated and lysed as in (E). Blue arrowhead indicates HOIL-1 Cter.

Next, we determined whether stimuli, that employ SHARPIN, modify its phosphorylation. Mimicking antigen receptor engagement with phorbol 12-myristate 13-acetate (PMA) plus ionomycin increased the level of SHARPIN phosphorylation, which was efficiently removed by lambda phosphatase (Fig. 1C-D). This was however not the case for all activating stimuli. The change in SHARPIN phosphorylation was induced by PMA, but not ionomycin alone, cell stress inducer Anisomycin or TNFα stimulation (Fig. 1E, EV1B). SHARPIN phosphorylation is therefore specifically increased upon antigen receptor engagement. By comparing the signaling pathways ignited by these stimuli, we noticed that ERK activation was correlated with stimulation-mediated SHARPIN phosphorylation (Fig. 1E, EV1B). Incubating cells with the MEK1/2 inhibitor Trametinib prior to antigen receptor engagement, via PMA plus ionomycin or CD3 plus CD28 stimulation, brought SHARPIN phosphorylation back to basal levels, but did not prevent this constitutive mark (Fig. 1F, EV1C). By contrast, inhibition of the NF-κB pathway with the PKC inhibitor Bisindolylmaleimide VIII (BIMVIII) had no effect (Fig. 1F, EV1C). To gain molecular insights, HOIP, SHARPIN and ERK1-FLAG were overexpressed in HEKT293T cells. Co-immunoprecipitation of FLAG-tagged ERK1 showed an interaction between SHARPIN and ERK1, which was lost in the absence of HOIP (Fig. EV1D). Likewise, SHARPIN phosphorylation could be induced by recombinant ERK1 when the LUBAC is isolated through pulled down of the HOIL-1 subunit (Fig. EV1E). This indicates that SHARPIN requires the LUBAC complex in order to be phosphorylated by ERK1. However, while incubating cell lysates with lambda phosphatase entirely removed SHARPIN phosphorylation, ERK1/2 inhibition only brings SHARPIN phosphorylation back to control conditions (Fig. 1A, 1F). Hence, it seems that at least two kinases are involved in the phosphorylation of SHARPIN, one, yet unidentified, that phosphorylates SHARPIN in resting conditions, and ERK1/2, which phosphorylates SHARPIN upon antigen receptor engagement.

### Serine 165 is the major phospho-acceptor site within SHARPIN

Mass spectrometry of SHARPIN, purified by SHARPIN immunoprecipitation of untreated Jurkat cells, was conducted to identify putative phosphorylation sites. In total, 20 peptides were recognized as SHARPIN, covering 45.5% of the protein sequence (Fig. EV2A). Five phosphorylated peptides were identified, and five phosphorylation sites were detected on S129, S131, S146, S165 and S312, with a probability of 65.20%, 50%, 92,44%, 100% and 89.49%, respectively (Fig. 2A, EV2B). Out of the 5 serines identified, S165 and S312 were the most conserved ones across species (Fig. 2B). To explore the details of SHARPIN phosphorylation, we engineered a stable Jurkat cell line deficient in *Sharpin* using the CRISPR/Cas9 technology (Fig. EV2C-D). Jurkat *Sharpin* knockout cells were subsequently rescued with an empty vector (EV), wild type (WT) SHARPIN, or phospho-dead mutants of putative SHARPIN phosphorylation sites (S165A, S312A, S165A+S312A (2SA)). S165 SHARPIN and 2SA SHARPIN mutants, in both unstimulated and stimulated conditions, had a faster migration in SDS-PAGE of SHARPIN than the WT or S312A SHARPIN (Fig. 2C, EV2E). This implicates S165 to be the main phosphorylation site of SHARPIN. Western blotting analysis with a phospho-specific S165 (p-S165) SHARPIN antibody, after immunoprecipitation of SHARPIN, confirmed that SHARPIN is constitutively phosphorylated and that this phosphorylation can be further induced upon PMA plus ionomycin activation while it remains unaffected by TNFα stimulation (Fig. 2D).

**Figure 2.**
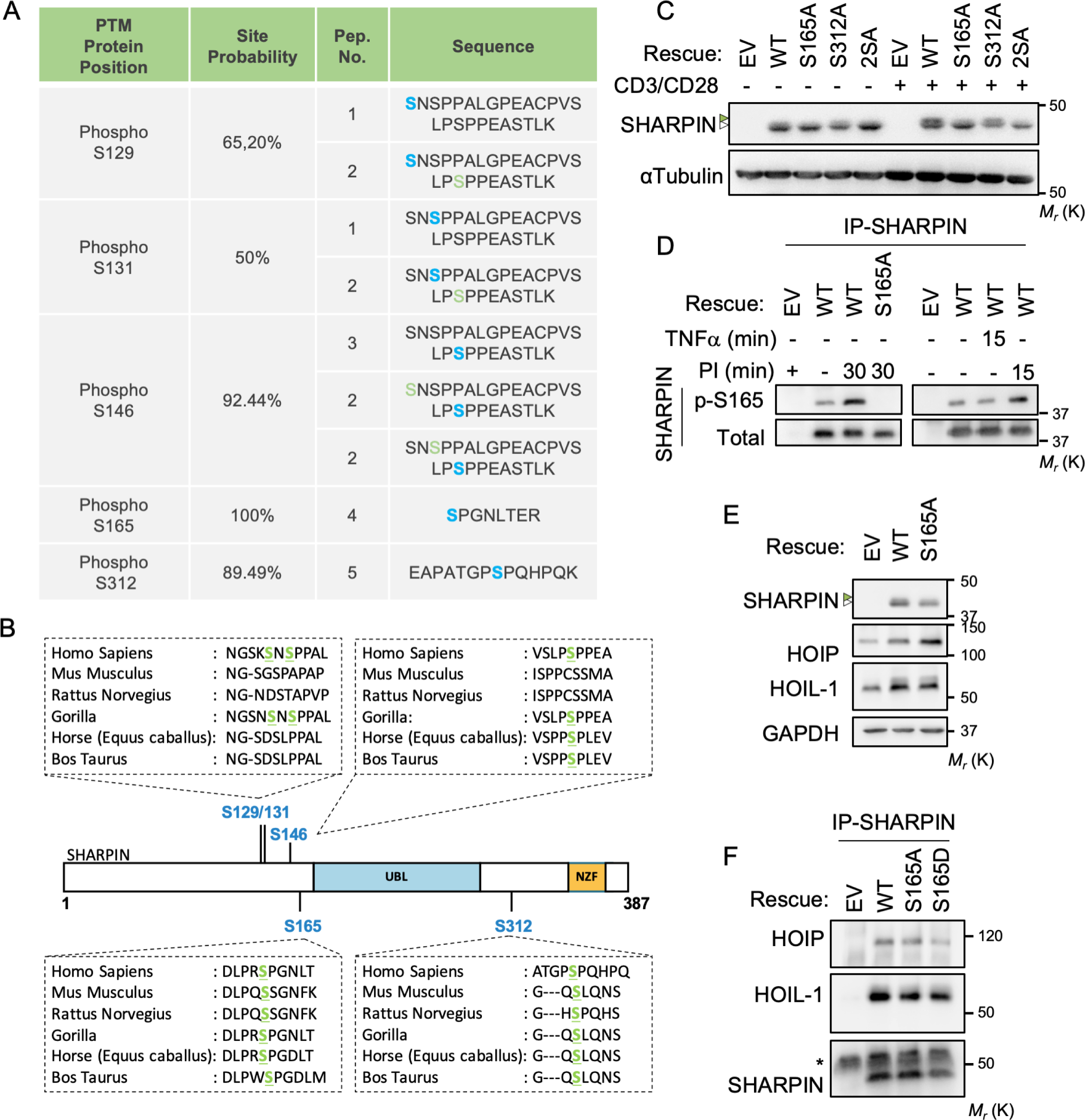
The Serine Residue of SHARPIN in Position 165 is a major Phosphorylation Site. A Mass spectrometry analysis of immunoprecipitates prepared from Jurkat cell lysates with antibodies to SHARPIN identified phosphorylation sites. The identified peptides were validated with an FDR < 1% and a minimum score of 30. The peptide score calculated in the Mascot search engine following -10log_10_(p), where p is the absolute probability. The position of the post-translational modification in the protein (PTM), the modification site probability calculated in Proline (site probability) the number of peptides identified, the peptide sequence with the identified amino acid in blue the localization and other amino acid modifications in green. B Sequence alignment of the amino acids surrounding the serine (S) 129, S131, S146, S165 and S312 residues. C Jurkat cells knockout for *Sharpin* were engineered by CRISPR/Cas9. Cells were complemented with an empty vector (EV), or with wild-type (WT), S165A-, S312A- or S165A plus S312A (2SA)-SHARPIN. Cells were stimulated for 30 min with antibodies to CD3 and CD28 (1 µg/mL each). Cell lysates were prepared and subjected to Western blotting analysis. White and green symbols show SHARPIN and phosphorylated SHARPIN, respectively. Molecular weight markers (*M*_*r*_) are indicated. D *Sharpin* knockout Jurkat cells reconstituted with EV, WT- or S165A-SHARPIN were treated with PI as in (C), or with 10 ng/mL TNFα. Cells lysates were Immunoprecipitated (IP) with antibodies specific to SHARPIN and subjected to Western blotting analysis with a mouse antibody specific for p-S165-SHARPIN. E Cell lysates from *Sharpin* knockout Jurkat cells reconstituted with an empty vector (EV), wild-type (WT-) or S165A-SHARPIN were subjected to Western blotting analysis with antibodies specific to the indicated antibodies. F Cell lysates from EV, WT-, S165A- or S165D-SHARPIN reconstituted *Sharpin* knockout Jurkat cells were submitted to SHARPIN immunoprecipitation (IP) prior to western blotting analysis for LUBAC components. * indicates the heavy chain of the SHARPIN antibody used for IP.

As previously reported [1–3,14,33], the stability of LUBAC is compromised in SHARPIN deficient cells, which results in a diminished NF-κB activation as observed by NF-κB reporter luciferase assay (Fig. EV2D, EV2F). Rescuing Jurkat *Sharpin* knockout cells with WT, S165A or phospho-mimetic S165D SHARPIN resulted in a replenishment of the LUBAC components HOIP and HOIL-1 (Fig. 2E), in a similar manner than the parental cells (Fig. EV2G). Hence, SHARPIN phosphorylation had seemingly no effect on the composition of the LUBAC (Fig. 2F).

### SHARPIN phosphorylation contributes to the optimal activation of NF-κB

The LUBAC is recruited to the TNFR1 complex I upon TNFα stimulation, where it is crucial for stabilizing the complex and subsequent NF-κB activation. It does so by modifying components of the TNFR1 complex I, including RIPK1, with linear ubiquitin (Methionine-1, M1) chains. This allows for a more efficient recruitment and retention of the IκB kinase (IKK) complex, consisting of IKKα, IKKβ and NEMO, with NEMO itself being modified with M1-chains. When activated, the IKK complex phosphorylates NF-κB inhibitors IκBs leading to their degradation. This liberates NF-κB to translocate into the nucleus where it can fulfill its transcription factor function [1–3,7,34]. Loss of LUBAC components leads to a switch from complex I to complex II, thereby inducing cell death by apoptosis or necroptosis [1,14,15]. We therefore studied the effect of SHARPIN phosphorylation on cell viability following TNFα stimulation. As previously shown [1–3], TNFα stimulation of *Sharpin* knockout cells reconstituted with an empty vector effectively reduced cell viability, while reconstitution with WT-, S165A-, or S165D-SHARPIN resulted in a similar protective effect against TNFα (Fig.3A). Even though phosphorylated SHARPIN is detected at the TNFR1, upon TNFα-FLAG stimulation (Fig. EV3A), recruitment of the phospho-dead SHARPIN to the TNFR1 complex I was normal, as was M1 ubiquitination at the TNFR1 receptor (Fig. EV3B). Likewise, TNFα stimulation did not drive any significant changes in total M1 ubiquitination or linear ubiquitination of the known substrate RIPK1 (Fig. 3B). Activation of immune cells by antigen receptor engagement leads to the assembly of a signaling complex of CARMA1, BCL10 and MALT1 (coined CBM complex). This complex serves as docking site for the LUBAC, which authorizes the recruitment and activation of IKK [17–19,33,35]. Within this signaling platform, the paracaspase MALT1 cleaves HOIL-1 [29–31]. We found that the phosphorylation status of SHARPIN did not influence the capability of MALT1 to cleave HOIL-1 upon antigen receptor engagement (Fig. 3C), implicating a normal assembly of the CBM complex. Furthermore, IκBα phosphorylation was unchanged in cells stimulated with TNFα or with PMA plus ionomycin (Fig. EV3C-D). Altogether; these results suggest that SHARPIN does not affect the recruitment of the LUBAC to the TNFR1 or CBM complexes, nor its ability to catalyze M1 chains and promote IKK activation.

**Figure 3:**
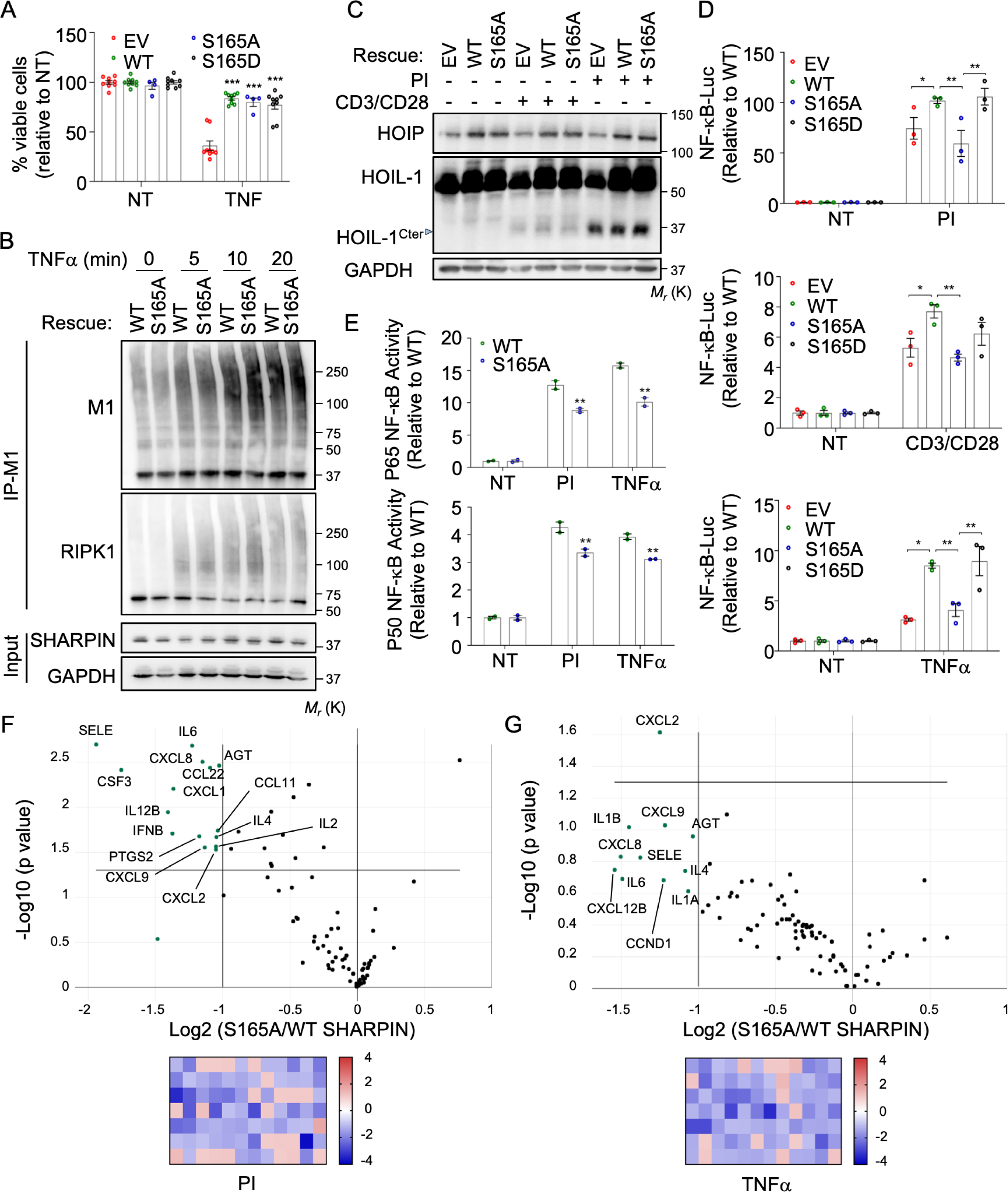
Serine 165 Phosphorylation of SHARPIN is crucial for the optimal Activation of NF-κB. A *Sharpin* knockout Jurkat cells expressing an empty vector (EV) or wild type (WT-), S165A- or S165D-SHARPIN were stimulated with 10 ng/mL TNFα for 24 h. Cell viability was assessed with CellTiter glo (mean ± SEM; ***P<0.001 by two-way ANOVA). B WT- and S165A-expressing Jurkat cells were stimulated with 10 ng/mL TNFα as indicated. Cell lysates were subjected to immunoprecipitation (IP) of linear ubiquitin (M1) prior to Western blotting analysis with the specified antibodies. Molecular weight markers (*M*_*r*_) are indicated. C *Sharpin* knockout Jurkat cells expressing an empty vector (EV) or wild type (WT-) or S165A-SHARPIN were stimulated with 20 ng/mL PMA plus 300 ng/mL ionomycin (PI) or CD3 plus CD28 (1 µg/mL each) for 30 min. Cell lysates were subjected to western blotting analysis. Blue symbol indicates HOIL-1 Cter cleavage band. D NF-κB reporter luciferase assay in the indicated Jurkat cells treated for 6 h with 20 ng/mL PMA plus 300 ng/mL ionomycin (PI), 1 µg/mL anti-CD3 plus 1 µg/mL anti- CD28, or with 1 ng/mL TNFα. Graph displays 1 out of 3 experiments done in triplicate (mean ± SEM; *p<0.05, **P<0.01 by two-way ANOVA). E WT- or S165A-SHARPIN reconstituted Jurkat cells were stimulated with PI or TNFα for 30 min as in (D) before performing TransAM NF-κB activation assay for p65 and p50 NF-κB subunits. The presented Graph show 1 out of 3 experiment performed in duplicate (mean ± SEM; *p<0.05, **P<0.01 by two-way ANOVA). F-G Volcano plot or RT^2^ profiler PCR array of human NF-κB signaling targets for cells stimulated for 4 h with PI (F) or TNFα (H). (G) Genes indicated on the volcano plot are significantly down regulated with a fold change > 2 in cells expressing S165A-SHARPIN when compared to WT-SHARPIN. On (G), CXCR2 is significantly down regulated with a fold change >2, while other genes indicated are borderline significant in S165A-SHARPIN expressing cells, compared to WT-SHARPIN. (F-G) Heatmap in the below volcano plot gives an overall view of the gene expression in the RT^2^ profiler PCR array.

Nonetheless, SHARPIN phosphorylation emerged as a key player in the optimal activation of NF-κB (Fig. 3D-G). NF-κB family members control the transcription of cytokines, and regulate cellular differentiation, survival and proliferation [36]. The NF-κB family is composed of five related transcription factors in mammals: p50, p52, p65 (also called RelA), c-Rel and RelB [37]. As expected, the transcription activity of NF-κB, as measured by a NF-κB reporter luciferase assay, was reduced in *Sharpin* knockout cells, reconstituted with an empty vector, stimulated through antigen receptor engagement or TNFα, when compared to cells expressing WT-SHARPIN (Fig. 3D). Interestingly, NF-κB activity was restored in cells expressing S165D-SHARPIN, but not S165A-SHARPIN (Fig. 3D), unveiling a key role for S165 phosphorylation. Likewise, cells expressing S165A-SHARPIN displayed a reduced DNA binding of the NF-κB subunits p65 and p50 upon antigen receptor engagement or TNFα stimulation (Fig. 3E). Of note, the binding of c-Rel was unchanged (Fig. EV3E), whereas p52 and RelB signals did not reach the threshold limits (data not shown). In keeping with these results, PCR array of human NF-κB signaling targets showed a significant diminished expression of numerous cytokines (CSF3, IL12B, IFNBA, IL6, CXCL9, CCL22, IL2 and IL4) and chemokines (CXLCL1, CXCL2, CXCL8, CCL11) in S165A-SHARPIN expressing cells treated with PMA and ionomycin (Fig. 3F, Fig. EV3F). Similar results were found when cells were stimulated with TNFα, albeit only the chemokine receptor CXCR2 reached statistical significance there is a trend towards reduced expression of other NF-κB signaling targets (Fig. 3G, EV3G). It should be mentioned that compared to PMA plus ionomycin stimulation, induction of NF-κB target genes by TNFα was weaker, which may also explain why differences upon TNFα stimulation are less striking. Nevertheless, we established that the phosphorylation of SHARPIN on S165 is participates in NF-κB upon antigen receptor engagement and TNFα stimulation.

In summary, we have discovered that a part of SHARPIN is constitutively phosphorylated in lymphoblastoid cells and identified S165 as the main phospho-acceptor residue. We also provide evidence that SHARPIN is further phosphorylated on the Serine 165 by ERK upon antigen receptor engagement, but not in response to TNFα (Fig. 1C). Yet, the constitutive phosphorylation of SHARPIN is pivotal for the optimal activation of NF-κB in response to antigen receptor engagement and TNFR ligation. This apparent dichotomy militates against a role for the inducible phosphorylation in NF-κB activation. It is therefore tempting to speculate that ERK-mediated phosphorylation of SHARPIN plays an independent function. Beside its crucial role in NF-κB signaling, a growing body of literature suggests that SHARPIN also acts as an inhibitor of the integrin adhesion receptors [38–40]. Pouwels *et al*. demonstrated that SHARPIN locates at and controls the detachment of cellular protrusions called uropods in lymphocytes, and that this is essential for lymphocyte movement [39]. As antigen receptor engagement delivers a stop signal to migrating T lymphocytes [41–43], ERK-dependent SHARPIN phosphorylation may have an impact on cell adhesion and migration. This description of at least two kinases targeting the same site, with different outcomes is reminiscent of what has been shown for MLKL (mixed lineage kinase domain-like pseudokinase). MLKL acute phosphorylation by RIPK3 at T357 and S358 triggers necrotic cell death [44], while basal phosphorylation on the same sites by a yet-to-be-defined kinase promotes the generation of small extracellular vesicles [45]. Lastly, the LUBAC may exist in different complexes, with different binding partners [46,47]. One could therefore speculate that different kinases could target different LUBAC complexes, and consequently exert selective functions.

Although necessary for complete NF-κB activity upon antigen receptor engagement and TNFα stimulation, how SHARPIN phosphorylation regulates the LUBAC functions and NF-κB signaling is unclear. The basal S165 SHARPIN phosphorylation appeared dispensable for TNFα-induced cell death, linear ubiquitination, assembly of CBM or TNFR1 complexes, and IKK activation. Additional work is therefore required to better understand this new layer of complexity in the regulation of NF-κB transcription activity. Nevertheless, this may also open up new avenues for therapeutic prospects of aggressive lymphoma, such as ABC DLBCL, for which the LUBAC and NF-κB activation is pivotal for survival [19,33]. *Hoip* and *Sharpin* deficiency in mice results in embryonic lethality or multiorgan inflammation, respectively, which is driven by aberrant TNFα-induced cell death [14,15,48–50]. Directly targeting SHARPIN phosphorylation for treatment may therefore circumvent the pitfall of inducing autoinflammatory diseases while targeting the NF-κB pathway specifically. The identification of the kinase responsible for constitutive SHARPIN phosphorylation will therefore be paramount to our future research.

## Material and Methods

### Cell culture

Jurkat E6.1 T lymphocytes, Hela, HEK293T and MCF7 cells were purchased from American Type Culture Collection (ATCC). U2932, SUDHL4, BJAB, RIVA, OCI-LY3, and OCI-LY19 cells were acquired from DSMZ. HBL1, HLY1 and TMD8 cells were kindly provided by Martin Dyer, Pierre Brousset, and Daniel Krappmann, respectively. Blood was obtained from healthy donors (Etablissement Français du Sang). Peripheral blood mononuclear cells (PBMC) were acquired by Ficoll density gradient. Primary CD4^+^ and CD8^+^ T cells were subsequentially isolated from PBMC using REAlease CD4 and CD8 microbeads kits, from Miltenyi as manufacturer’s instructions. Cell viability was assessed using CellTiter-Glo following manufacturer’s instructions (Promega) following stimulation with 10 ng/mL TNFα (R&D systems) for 24h.

### Immunoblotting

Jurkat E6.1, PBMC or primary T cells were stimulated with 20 ng/mL Phorbol 12-Myristate 13 Acetate (PMA, Merck), 300 ng/mL ionomycin (Merck), PMA plus ionomycin, 1 μg/mL CD3 plus 1 μg/mL CD28 (Becton Dickinson Biosciences), 12.5 mg/mL anisomycin (Merck), 10 ng/mL TNFα (R&D systems) or TNFα-Flag (Enzo Life Sciences). To ensure inhibition of certain signaling pathways Jurkat E6.1, PBMC or primary T cells were pre-treated during 1h with 1 μM Trametinib (Selleckchem), 500 nM Bisindolylmaleimide VIII (BIMVIII, Enzo), 50 μM SP6000125 (Cell Signaling), or 5 μM SB203580 (Selleckchem). Cells were washed with ice-cold PBS and subsequently lysed using two possible lysis buffers, as indicated throughout the manuscript. (1) Cells pellets were incubated with 188 μl of Buffer A (10 mM HEPES pH7.9, 10 mM KCL, 0.1 mM EDTA, 0.1 mM EGTA, 1mM DTT and 1 mM Na_3_VO_4_) for 5 min on ice. 12 μl of Buffer A containing 10% Igepal was added for another 5 min on ice and samples were subsequently centrifuged at 1,000 g for 3 min. Supernatant was collected. (2) Cell pellets were lysed with TNT buffer (50 mM Tris-HCl (pH 7.4), 150 mM NaCl, 1% Triton X-100, 1% Igepal, 2 mM EDTA), and incubated for 30 min on ice. Extracts were cleared by centrifugation at >10 000 x g. Lysis buffers were supplemented with 1x Halt Protease Inhibitor cocktail (ThermoFisher Scientific). Protein concentration was determined using the Micro BCA Protein Assay kit (ThermoFisher scientific), for both lysis buffers. Lambda phosphatase experiments used either Buffer A or TNT lysis without EDTA or EGTA for cell lysis. 10 μg of protein was incubated with 600 Units of lambda phosphatase, 1x NEB buffer for Protein Metallophosphatase and 1mM MnCl_2_ (New England Biolabs) for 30 min at 30°C. 10 µg of proteins was incubated with 2X Laemmli buffer (Life Technologies) at 95°C for 3 min, separated by SDS-PAGE using a 3-15% Tris-Acetate gel and transferred to nitrocellulose membranes (GE Healthcare). Densitometry analysis was performed by the Image J software (National Institutes of Health).

The following antibodies were used for analysis by immunoblot: SHARPIN (A303-559A), HOIP (A303-560A), and IκBβ (A301-828A) antibodies were purchased from Bethyl. HOIL-1 (sc-393754), αTubulin (sc-8035), CYLD (sc-137139) and GAPDH (sc-32233) antibodies were obtained from Santa Cruz Biotechnology. Phospho-ERK (#9106), IκBα (#9242), phospho-IκBα (#9246), phospho-p38 (#9215), phospho-JNK (#9255) and RIP1 (#3493) were acquired from Cell Signaling Technologies. Anti-linear Ubiquitin (MABS199) was procured from Millipore. For detection of the phospho-S165 from of human SHARPIN, five Balb/C mice were primed via intraperitoneal injection with complete Freund adjuvant and boosted two times at two-week intervals with incomplete Freund adjuvant with a synthetic phosphorylated peptide (DLPR(Sp)PGNLTERC) conjugated to KLH. Mouse blood was collected from the submandibular vein and serum reactivity was confirmed by ELISA with phosphorylated and non-phosphorylated biotinylated peptide. Mouse serum was used to reveal phospho-S165 SHARPIN.

### Immunoprecipitation (IP) and ERK recombinant kinase assay

Cells were washed with ice-cold PBS, lysed with TNT buffer and incubated for 30 min on ice. Extracts were cleared by centrifugation at >10,000 g. Protein concentration was determined using the Micro BCA protein kit. Samples were precleared with Protein G Sepharose (Merck) for 30 min, and subsequently incubated with 1 μg of antibody and protein G Sepharose for a duration of 2h. Flag pull down was performed incubating cleared cell lysates with Anti-flag M2 affinity gel (Merck) during 2h. SHARPIN (A303-559A, Bethyl), HOIL-1 ((sc-393754, Santa Cruz Biotechnology) and linear ubiquitin, kindly provided by VM Dixit (Genentech), antibodies were used for IP. ERK1/2 kinase assay was performed from beads immunoprecipitated for HOIL-1. Beads were washed in kinase buffer (5 mM MOPS pH 7.2, 5 mM MgCl_2_, 1 mM EGTA, 0.4 mM EDTA, 0.05 mM DTT). Beads were subsequently incubated with kinase buffer, 2mM ATP, 350 ng/mL active untagged ERK1 (Merck), 350 ng/mL active ERK2-GST tagged (Merck) or 175 ng/mL ERK1 plus 175 ng/mL ERK2-GST for 30 min at 30°C.

### Mass Spectrometry

Jurkat E6.1 cells were lysed in TNT lysis buffer supplemented with 1x Halt protease inhibitor cocktail. Immunoprecipitation of SHARPIN was performed as described above. Proteins were separated by SDS-PAGE. The Colloidal Blue Staining kit (LC6025, Invitrogen) was used to stain protein as per manufacturer’s instructions. Gel was excised between 37 and 50 kDa, to ensure the presence of SHARPIN, and subsequently cut into smaller fragments. Fragments were washed alternating between 100 mM Ammonium Bicarbonate and Acetonitrile. Disulfide bonds of proteins were reduced, with using 65 mM DTT for 15 min at 37°C, and subsequently alkylated by 135 mM Iodoacetamide for 15 min at room temperature. Enzymatic digestion of proteins was performed at 37°C overnight by incubating the gel pieces in 25 ng/mL pH 8.5 trypsin solution (Sequenced Grade Modified Trypsin, ref V511A, Porcine, Promega). Digested peptides were extracted by incubation in 70% and 100% acetonitrile for 20 min each. Peptide extracts were evaporated and afterward reconstituted in 0.1% formic acid. Peptide extracts were analyzed by liquid nano-chromatography (nanoLC) nanoElute coupled with a TimsTOF Pro mass spectrometer (Bruker), as previous described by [51]. Generated peaks were analyzed with the Mascot database search engine (MatrixScience version 2.5.01) for peptide and protein identification. The peaks were queried simultaneously in the: UniprotKB Human (release 20191016, 20656 sequences) database and a decoy database that was interrogated in parallel to estimate the number of false identifications and to calculate the threshold at which the scores of the identified peptides are valid. Mass tolerance for MS and MS/MS was set at 15 ppm and 0.05 Da, respectively. The enzyme selectivity was set to full trypsin with one miscleavage allowed.

Protein modifications were fixed carbamidomethylation of cysteines, variable oxidation of methionine, variable phosphorylation of serine, threonine or tyrosine. Identified proteins are validated with an FDR < 1% at PSM level and a peptide minimum score of 30, using Proline v2.0 software [52]. Proteins identified with the same set of peptides are automatically grouped together. Analysis of post-translational modifications was also executed using Proline v2.0 Software. Localization confidence values for peptide ions were exported from Mascot [53] and a site probability is calculated for a particular modification according to the number of peptides confidently detected with this site modification.

### CRISPR/Cas9 knock out and SHARPIN reconstituted cell lines

LentiGuide and LentiCRISPRv2 vectors (GeCKO, ZhangLab), were cloned to contain SHARPIN guide RNA (5’ CACCGTGGCTGTGCACGCCGCGGTG 3’), as described before (Sanjana et al., 2014; Shalem et al., 2014). LentiCRISPRv2, together with the packaging vectors PAX2 and VSV-G, were transfected in HEK293T cells using a standard calcium phosphate protocol, as previously published [54]. Supernatant, containing the lentiviral particles, was collected after 48 h, and used to infect 10^6^ Jurkat cells in presence of 8 μg/mL Polybrene (Santa Cruz Biotechnology). Jurkat cells expressing the LentiCRISPRv2 vector were selected by adding 1μg/mL Puromycin to their media, and subsequently dilution cloned. Single-cell clones were then picked and tested for functional Cas9 cutting of SHARPIN (Fig.EV2C). SHARPIN was cloned from a pCMV3flag9SHARPIN vector (Addgene), into a pCMH-MSCV-EF1a-puroCopGFP vector (SBI). Site directed mutagenesis was performed using the pCMH-MSCV-EF1a-puroCopGFP vector containing wild type (WT) SHARPIN, to substitute the serine (S) residues on 165 or/and 312 to an alanine (A) or an aspartic acid (D). CRISPR/Cas9 resistance was achieved by site directed mutagenesis of the PAM sequence on site 29, substituting an Arginine (AGG) to an Arginine (AGA).

### NF-κB assays

NF-κB luciferase assay was performed as previously described [20,29]. DNA binding of the NF-κB subunits was measured using TransAM NF-κB activation assay (Actif Motif), as per manufacturer’s instructions, with reconstituted Jurkat cells were treated with 20 ng/mL PMA plus 300 ng/mL Ionomycin or 10 ng/mL TNFα during 30 min. For the RT^2^ profiler PCR array of human NF-κB signaling targets, cells were treated with 20 ng/mL PMA plus 300 ng/mL Ionomycin or 10 ng/mL TNFα for 4h. RNA was extracted using the Nucleospin RNAplus kit (Macherey-Nagel), following manufacturer’s instructions. 2 mg of RNA was reverse transcribed using the Maxima Reverse Transcriptase kit (ThermoFisher). RT^2^ profiler PCR array of human NF-κB signaling targets was performed as instructed by the manufacturer (Qiagen).

### Statistical analysis

Statistical analysis, comparing multiple groups was performed using two-way ANOVA on rank test with Tuckey’s post hoc test in GraphPad Prism 7 software. TransAM NF-κB activation assays were analyzed using a 2-way ANOVA test with a Sidak correction for multiple analysis. RT^2^ profiler PCR array was performed using the online software provided by the manufacturer (https://dataanalysis2.qiagen.com/pcr). In short, the p-values were calculated based on a student t-test of the replicate 2^(-DC_T_) values for each gene in the control group and treatment group. The p-value calculation used is based on a parametric, unpaired, two sample equal variance, two tailed distribution. P-values < 0.05 were considered as significant.

## Acknowledgments

This research was funded by an International Program for Scientific Cooperation (PICS, CNRS), Fondation pour la Recherche Médicale (Equipe labellisée DEQ20180339184), Fondation ARC contre le Cancer (NB), Ligue nationale contre le cancer comités de Loire-Atlantique, Maine et Loire, Vendée (JG, NB), Région Pays de la Loire et Nantes Métropole under Connect Talent Grant (JG), the National Research Agency under the Programme d’Investissement d’Avenir (ANR-16-IDEX-0007), and the SIRIC ILIAD (INCa-DGOS-Inserm_12558). AT and SR hold post-doctoral fellowships from la Fondation ARC contre le Cancer. TD is a PhD fellow funded by Nantes Métropole. This work was also supported by grants from Biogenouest, Infrastructures en Biologie Santé et Agronomie (IBiSA) and Conseil Régional de Bretagne awarded to CP.

## Author contribution

An Thys: Conceptualization, Methodology, Formal analysis, Investigation, Writing – original draft, Visualization, Funding acquisition. Kilian Trillet, Sara Rosinska, Audrey Gayraud, Tiphaine Douanne, Clotilde Renaud, Luc Antigny and Régis Lavigne: Investigation. Yannic Danger and Franck Vérité: Resources. Emmanuelle Com: Methodology, Formal analysis, Investigation. Charles Pineau: Methodology. Julie Gavard: Writing – Review and Editing, Supervision, Funding acquisition. Nicolas Bidère: Conceptualization, Methodology, Investigation, Writing – original draft, Visualization, Funding acquisition

## Conflict of interest

The authors declare that they have no conflict of interest.

## Expanded View Figure legends

**Figure EV1, relative to Figure 1.**
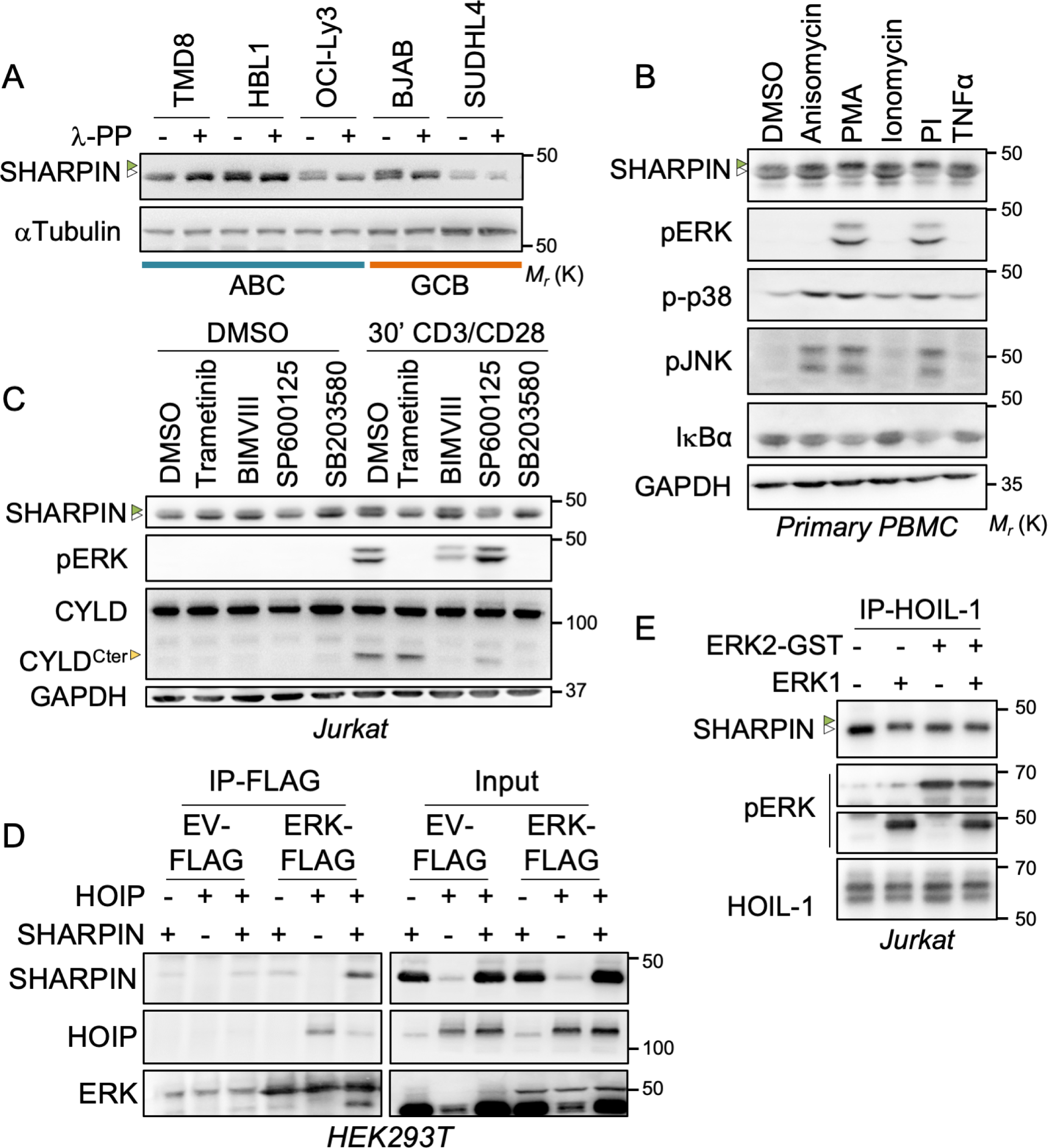
SHARPIN Phosphorylation by ERK upon Antigen Receptor Engagement. A Western blotting analysis of a panel of patient-derived diffuse large B-cell lymphoma (DLBCL) cell lines. Cell lysates were treated with lambda phosphatase (λPP) as indicated, and subjected to Western blotting analysis. Green and white arrowheads indicate phosphorylated and unphosphorylated SHARPIN, respectively. Molecular weight markers (*M*_*r*_) are indicated. B Primary peripheral blood mononuclear cells (PBMC) were stimulated with 12.5 µg/mL Anisomycin, 20 ng/mL PMA, 300 ng/mL ionomycin, PMA plus ionomycin or 10 ng/mL TNFα for 30 min. Cell lysates were subjected to Western blotting analysis. C Jurkat cells were pretreated with 1 µM Trametinib (MEK1/2 inhibitor), 500 nM Bisindolylmaleimide VIII (BIMVIII) (PKC inhibitor), 1 µM, 50 µM SP6000125 (JNK inhibitor), or 5 µM SB203580 (p38 inhibitor), and subsequently treated with CD3 plus CD28 (1μg/μL each). Western blotting analysis was performed as indicated. Treatment with SB203580 gave non-specific results as it also regulated ERK phosphorylation. Yellow symbol indicates CYLD Cter cleavage band. D HEK293T cells were co-transfected with plasmids encoding for HOIP, SHARPIN or HOIP plus SHARPIN together with ERK1-FLAG. Cells were lysed with TNT. Cell lysates were immunoprecipitated using anti-FLAG M2 affinity gel and subjected to Western blotting analysis. E The LUBAC was pulled down by immunoprecipitation of Jurkat cell lysates using the HOIL-1 antibody. *In vitro* kinase assay was performed by adding active ERK1, ERK2-GST or ERK1 plus ERK2-GST recombinant proteins to HOIL-1 pulled down beads. Western blotting analysis was performed for the indicated antibodies. White and green arrows show SHARPIN and phosphorylated SHARPIN, respectively. Molecular weight markers (M_r_) are indicated.

**Figure EV2, relative to Figure 2.**
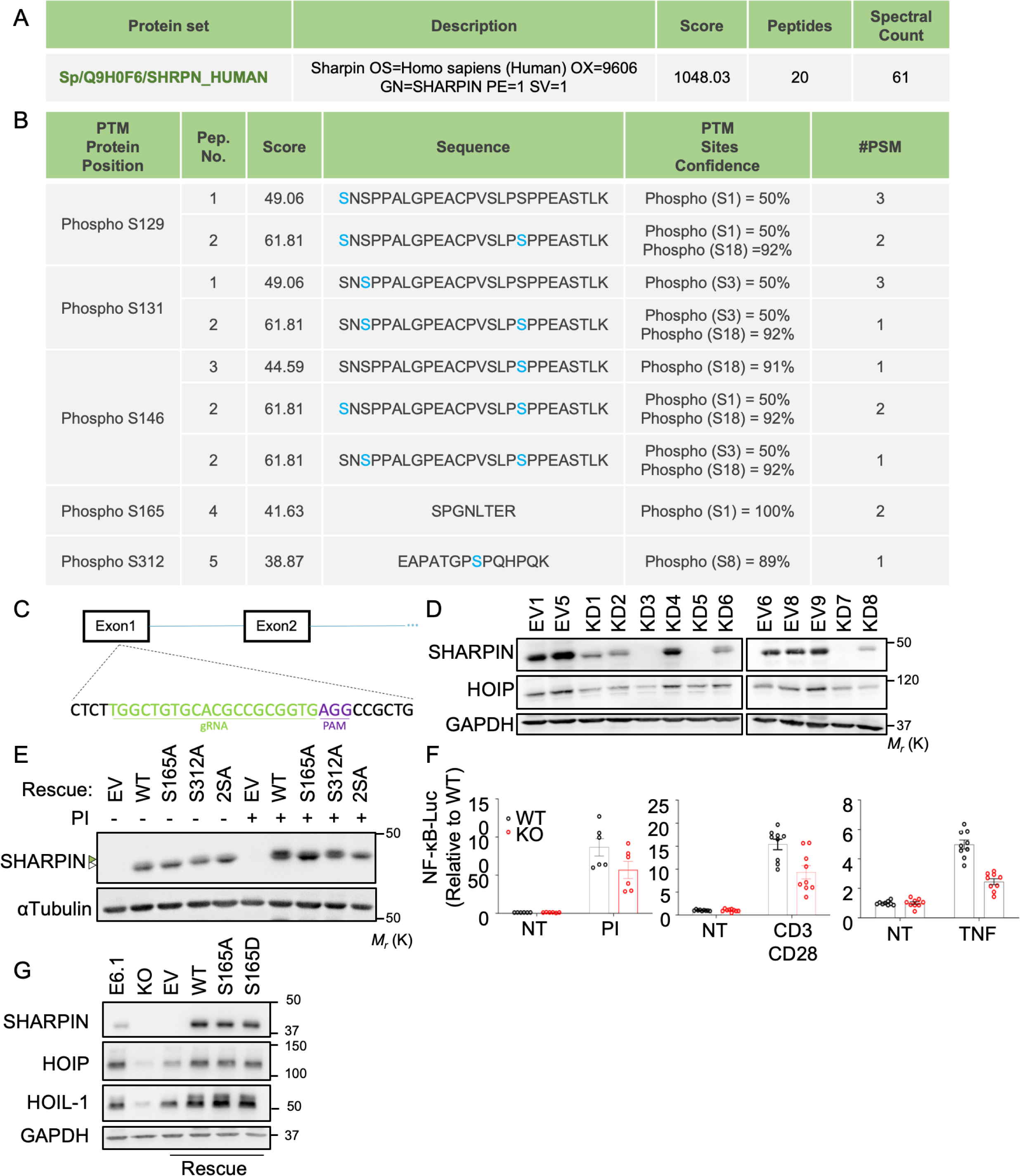
Characterization of SHARPIN knockout Cells. A SHARPIN identified during mass spectrometry analysis (peptide rank1, peptide score > 30, FDR<1% at PSM level). The protein score is the sum of the unique peptide score calculated following -10log_10_(p), where p is the absolute probability. Peptide count and spectral count are indicated. B Mass spectrometry analysis of immunoprecipitates prepared from Jurkat cell lysates with antibodies to SHARPIN identified phosphorylation sites. The identified peptides were validated with an FDR < 1% and a minimum score of 30. The peptide score calculated in the Mascot search engine following -10log_10_(p), where p is the absolute probability. The position of the post-translational modification in the protein (PTM), the number of peptides identified, the peptide score, the peptide sequence with the modified amino acid in blue the localization confidence value of the post-translational modification calculated in the Mascot search engine (ptm sites confidence) and the number of spectra detected for each peptide (#PSM or peptide-spectrum match). C Scheme showing the sgRNA target and PAM site on SHARPIN for the LentiCRISPR vector containing SHARPIN sgRNA. D Western blotting analysis of Jurkat single cell clones targeted with sgSHARPIN LentCRISPRv2 (knock down, KD or an empty vector (EV). E Jurkat *Sharpin* KO cells were complemented with an empty vector (EV), or with wild-type (WT), S165A-, S312A- or S165A+S312A (2SA)-SHARPIN. Cells were stimulated for 30 min with 20 ng/mL PMA plus 300 ng/mL ionomycin. Cell lysates were subjected to Western blotting analysis. White and green symbols show SHARPIN and phosphorylated SHARPIN, respectively. Molecular weight markers (*M*_*r*_) are indicated. F NF-κB reporter luciferase assay of Parental Jurkat and Jurkat knockout (KO) cells treated for 6 h with 20 ng/mL PMA plus 300 ng/mL ionomycin, CD3 plus CD28 (1 µg/mL each) or 1 ng/mL TNFα (mean ± SEM; *p<0.05, **P<0.01 by one-way ANOVA, n=2). G Cell lysates from Jurkat, *Sharpin* KO Jurkat cells or cells reconstituted with EV, WT-, S165A- or S165D-SHARPIN. Western blotting analysis was performed for the indicated proteins.

**Figure EV3, relative to Figure 3.**
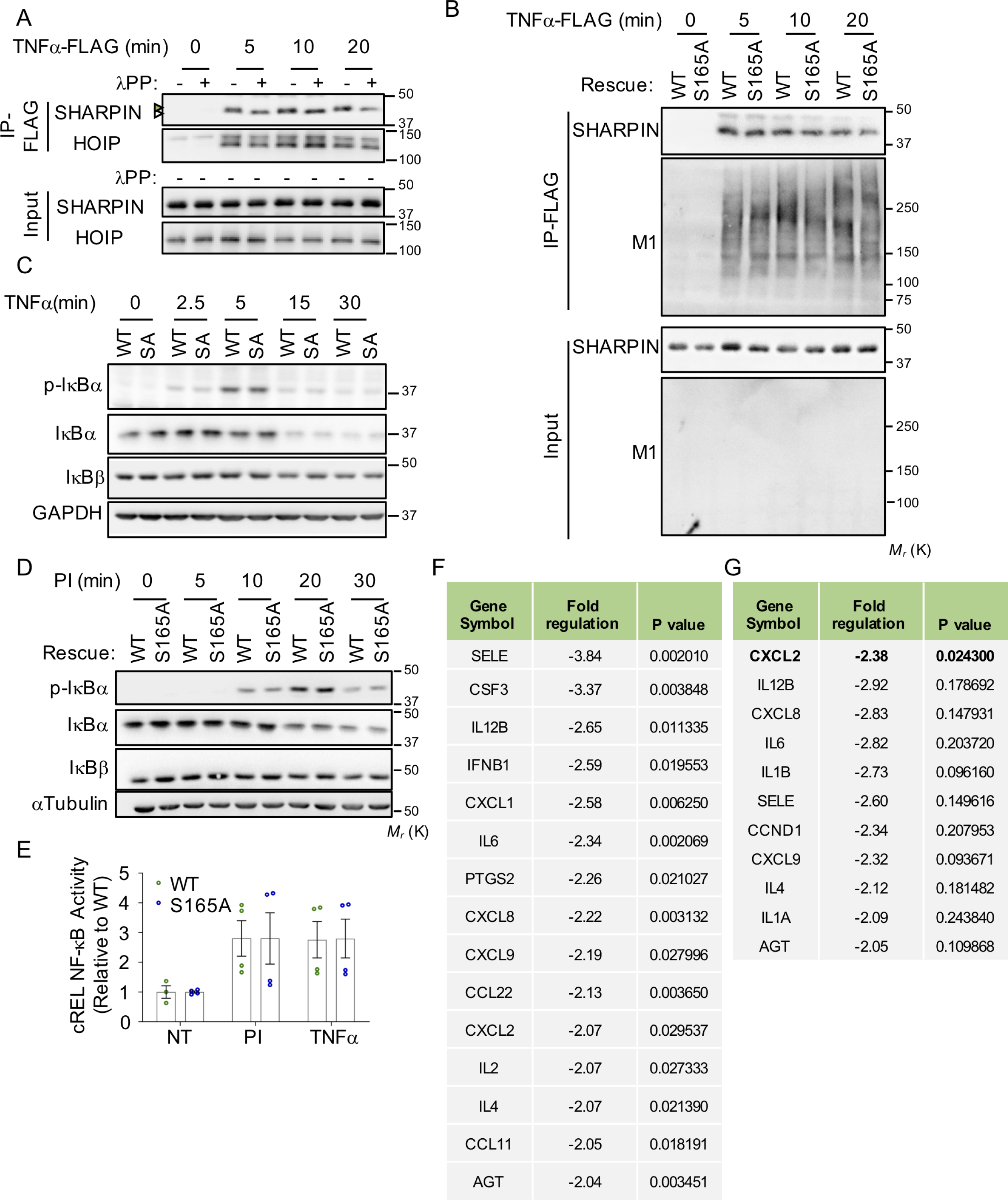
A-B Jurkat cells (A) or *Sharpin* knockout (KO) Jurkat cells, reconstituted with wild type (WT-) or S165A-SHARPIN (B) were stimulated with 100 ng/mL TNFα-FLAG for the indicated time points.). Cell lysates were immunoprecipitated using anti-FLAG M2 affinity gel (A-B). Beads were incubated with or without lambda phosphatase (λPP) (A) and subjected to Western blotting analysis (A-B). White and green symbols show SHARPIN and phosphorylated SHARPIN, respectively. Molecular weight markers (*M*_*r*_) are indicated. C-D Cell lysates of WT- and S165A-expressing Jurkat were treated with 10 ng/mL TNFα (C) or with 20 ng/mL PMA plus 300 ng/mL ionomycin (PI) (D) as indicated and subjected to Western blotting analysis. E WT- or S165A-SHARPIN reconstituted Jurkat cells were stimulated with PI or TNFα for 30 min as in (B-C) before performing TransAM NF-κB activation assay for cREL NF-κB subunit. (mean ± SEM; *p<0.05, **P<0.01 by two-way ANOVA, n=2). F-G List of RT^2^ profiler PCR array of human NF-κB signaling targets with an expression fold change >2 in in cells expressing S165A-SHARPIN when compared to WT-SHARPIN. Cells were treated during 4h with 20 ng/mL PMA plus 300 ng/mL ionomycin (F) or 10 ng/mL TNFα (G). Fold regulation and p-value are listed.

